# Allocation pattern of fruiting bodies in plasmodial slime molds, and threshold size for sporulation of *Physarum polycephalum*

**DOI:** 10.64898/2026.07.06.736535

**Authors:** Sota Takahashi, Yukinori Nishigami, Atsushi Taniguchi, Toshiyuki Nakagaki

**Affiliations:** Graduate School of Life Science, Hokkaido University, N10W8, Sapporo, 0600810, Hokkaido, Japan.; Research Institute of Electronic Science, Hokkaido University, N20W8, Sapporo, 0010020, Hokkaido, Japan.

**Keywords:** habitat selection, sporulation, single cell organism, plasmodium, *Physarum polycephalum*, survival, protists

## Abstract

The plasmodium of Myxogastria (a group of amoeboid protists) species often crawls around the forest floor to feed while searching for places to form fruiting bodies for reproduction (sporulation). Certain environmental factors that trigger sporulation have been reported; however, other unknown factors are also expected. In this study, we reported field observations of *Physarum rigidum* and *Fuligo septica*. Inspired by the field observation, we examined the effects of multiple factors on sporulation in laboratory experiments using *Physarum polycephalum*. We found that:(1) there was a critical body size below which sporulation did not occur under our experimental conditions and (2) the plasmodium selected its sporulation sites from the available landscape of the experimental arena: dry and low sites for the majority and dry and high sites for the minority. Further analysis revealed that they preferred the edge area at the high site. We discuss the possible ecological importance of the threshold and location preference.

## 1 Introduction

Slime molds belonging to the protist group are ecologically important because they are widespread predators of bacteria living on the forest floor and crucial prey for large insects (Tiunov et al. 2015; Potapov et al. 2022; Sugiura et al. 2019) owing to the multinucleate plasmodium life cycle stage that resembles a primitive multicellular organization. Plasmodia weighing several grams are typically found in the field and crawl at 1 cm/h around complex, uneven outdoor environments while searching for a suitable location to form many fruiting bodies (FBs; spore formation or sporulation for reproduction). The transformation from motile plasmodia to sessile FBs has been studied because of the drastic morphogenesis process, which is completed in only one day (Constantineanu 1906; Gray 1936; Sauer et al. 1969; Poetsch et al. 1989; Clark and Haskins 2015; Werthmann and Marwan 2017; Glöckner and Marwan 2017).

In *Physarum polycephalum*, numerous studies have identified the key factors that initiate sporulation and subsequent intracellular metabolic reactions and gene regulation. In particular, a combination of starvation (Camp 1937; Chapman and Coote 1982) and light (Gray 1938) is essential for inducing sporulation. This notion is supported by studies on starvation-related gene expression (Sauer et al. 1969; Werthmann and Marwan 2017; Glöckner and Marwan 2017) and metabolic production (Wilkins and Reynolds 1979; Renzel et al. 2000; Akitaya et al. 1985), with action spectra from ultraviolet to infrared light (Starostzik and Marwan 1995b; Nakagaki et al. 1996) identified in starved plasmodium.

In the field, plasmodium from species such as *Physarum, Fuligo, Stemonitis* crawls across different microniches on decaying logs, including dark humid places at the bottom of logs and sunny dry places at the top (Stahl 1884; Lubbock 1892; Gray 1936). Although such niches are often rich in food, FBs are found in various niches at the bottom and top of logs. Each species has characteristic morphologies, numbers, and allocation patterns of FBs that have evolved under selective pressure (Schnittler 2001; Schnittler and Tesmer 2008). Different species choose different types of niches (Vlasenko et al. 2018). Hence, there may be unknown factors at work, including internal conditions of the slime mold itself, as well as external conditions such as undulation and elevation of the ground it crawls upon.

To obtain a qualitative hypothesis on these factors, we began with field observations. We observed two species (*Physarum rigidum* and *Fuligo septica*) that were encountered by chance in the field. The field observation was limited to a single time. Experimental methods for FB formation have not been established in slime molds except for a few species. Hence, it was impossible to observe the FB formation of the two species in the laboratory.

Motivated by these field observations, we investigated the possible effects of factors such as body size, substratum modulations, and elevation within the habitat landscape on sporulation in *P. polycephalum*. To shed light on habitat selection for sporulation, we observed how plasmodia explored various available niches in an experimental arena and which areas were used for sporulation.

Laboratory experiments showed that plasmodia smaller than a certain threshold did not crawl on a dry surface and sporulate simultaneously. Additionally, after exploring the various niches, plasmodia allocated FBs to two different places: low places for most FBs and high places for a few. In particular, FBs were concentrated at the edge area of the high place.

The ecological importance of the size threshold and location preference is discussed in the Discussion. Finally, we discuss the potential for future ethological research on sporulation in slime molds.

## 2 Materials and Methods

### 2.1 Field observation and recording of the behavior

#### 2.1.1 Crawling trajectory and locations of FB formation in

##### Physarum rigidum

The sporulation process of a single plasmodium of *Physarum rigidum* was recorded under natural conditions (Fig. 1) from 16:00 on June 28 to 10:00 on July 5, 2025 at the Sapporo Campus of Hokkaido University (Sapporo, Hokkaido, Japan).

**Fig. 1.**
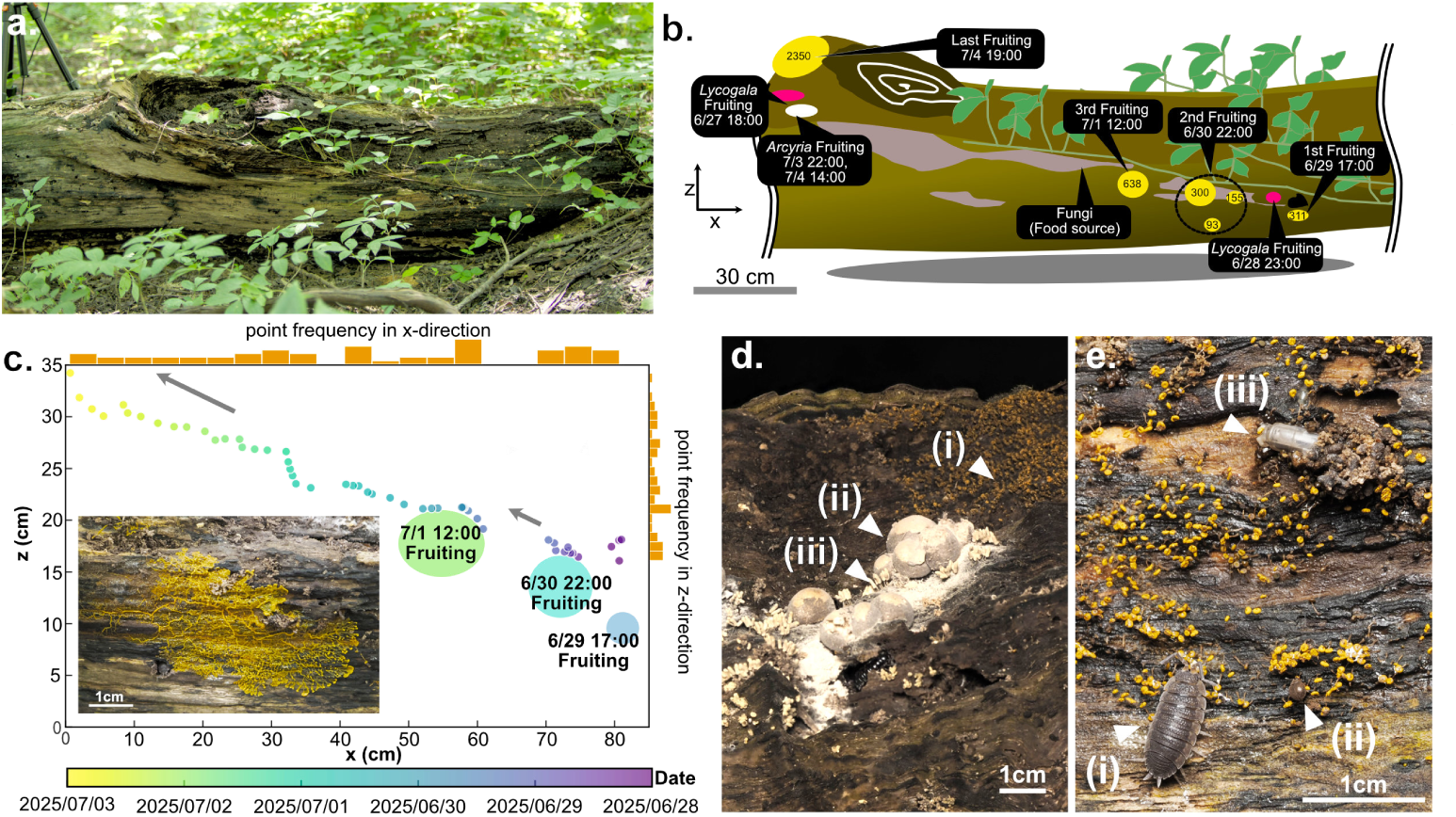
Behavior of *Physarum rigidum* plasmodium in the field. **a**. Lateral view of observed rotten log recorded at 9:12 on July 5. **b**. Schematic illustration for the crawling trajectory and sporulation, transcribed via time-lapse imaging. The number on the yellow circles indicates the number of fruiting bodies (FBs) formed at that point. **c**. The time course of position of the plasmodium (the position of the top leftmost tip of plasmodium), plotted every 2 h from June 28 to July 2. The origin of the *x* axis was determined by the initial position of the plasmodium. The origin of the *z* axis is the surface of the ground. The color of the circle symbols indicates the time and date according to the color chart shown at the bottom of the graph (violet at the beginning and yellow at the end of the observation period). The plasmodium migrated from the lower right on June 28, 2025 to the upper left on July 3, 2025 over six days while initiating FB formation three times, at 17:00 on June 29, at 22:00 on June 30, and at 12:00 on July 1. We counted the number of the data symbols along *x−* and *z−*axes and indicated them by the histograms (painted by orange color) on the top and right, respectively, of the graph panel. At the two positions, (*x, z*) = (70, 15) and (55, 20), as the column length on the top of graph and bar length on the left were longer, the plasmodium stayed at these positions for a long time before FB formation. Inset: A part of the crawling front captured at 13:00 on July 1. **d, e**. Interaction with other species. **d**. The topmost part of the tree photographed at 16:24 on July 6. The remaining part of the *Physarum* plasmodium reached the highest part of the tree and sporulated (i), neighboring the FBs of *Lycogala epidendrum* (ii) and *Arcyria cinerea* (iii). **e** Scene around the third fruiting patch photographed at 11:09 on July 2. Three arthropods visited the patch: (i) a species of woodlouse, (ii) *Aspidiphorus sp.*, and (iii) a fly larva.

Capturing the crawling trajectory: The front of the crawling organism was photographed every 10 min using a digital camera with an LED ring-light attachment (Tough TG-7 and LG-1; OM Digital Solutions, Japan) mounted on a tripod. The LED was illuminated only during autofocusing and exposure to light. The camera position was manually adjusted as the organism moved out of the field of view. After recording the entire process, the surrounding environment was photographed, including the entire trajectory of crawling.

Sporulation timing and crawling trajectory were determined from the recorded images. The onset of sporulation was estimated based on morphological changes in the plasmodium as seen in the recorded images and confirmed in situ by close-up imaging of FBs with a digital camera (OM-1 Mark II and M.ZUIKO DIGITAL ED 90mm F3.5 Macro IS PRO, OM Digital Solutions, Japan) and a flashlight (v350, Godox Photo Equipment, China). The number of FBs was counted in close-up images.

All researches were conducted in accordance with the laws and regulations of Japan and Hokkaido University.

#### 2.1.2 Climbing behavior before FB formation in *Fuligo septica*

We conducted field observations of climbing behavior before sporulation in *Fuligo septica* on a fallen tree in July 2024 at the Sapporo Campus of Hokkaido University (Sapporo, Hokkaido, Japan; Fig. 2). Climbing behavior was recorded using a digital camera (ILCE-7SM2, Sony, Japan) with a macro lens (EF 100mm L macro, Canon, Japan). The camera was set to record in time-lapse mode; images were acquired every 10 min from 18:00 on July 8 to 10:00 on July 9, 2024. The exposure, color temperature, white balance, and noise reduction level of each image were independently adjusted for visibility.

**Fig. 2.**
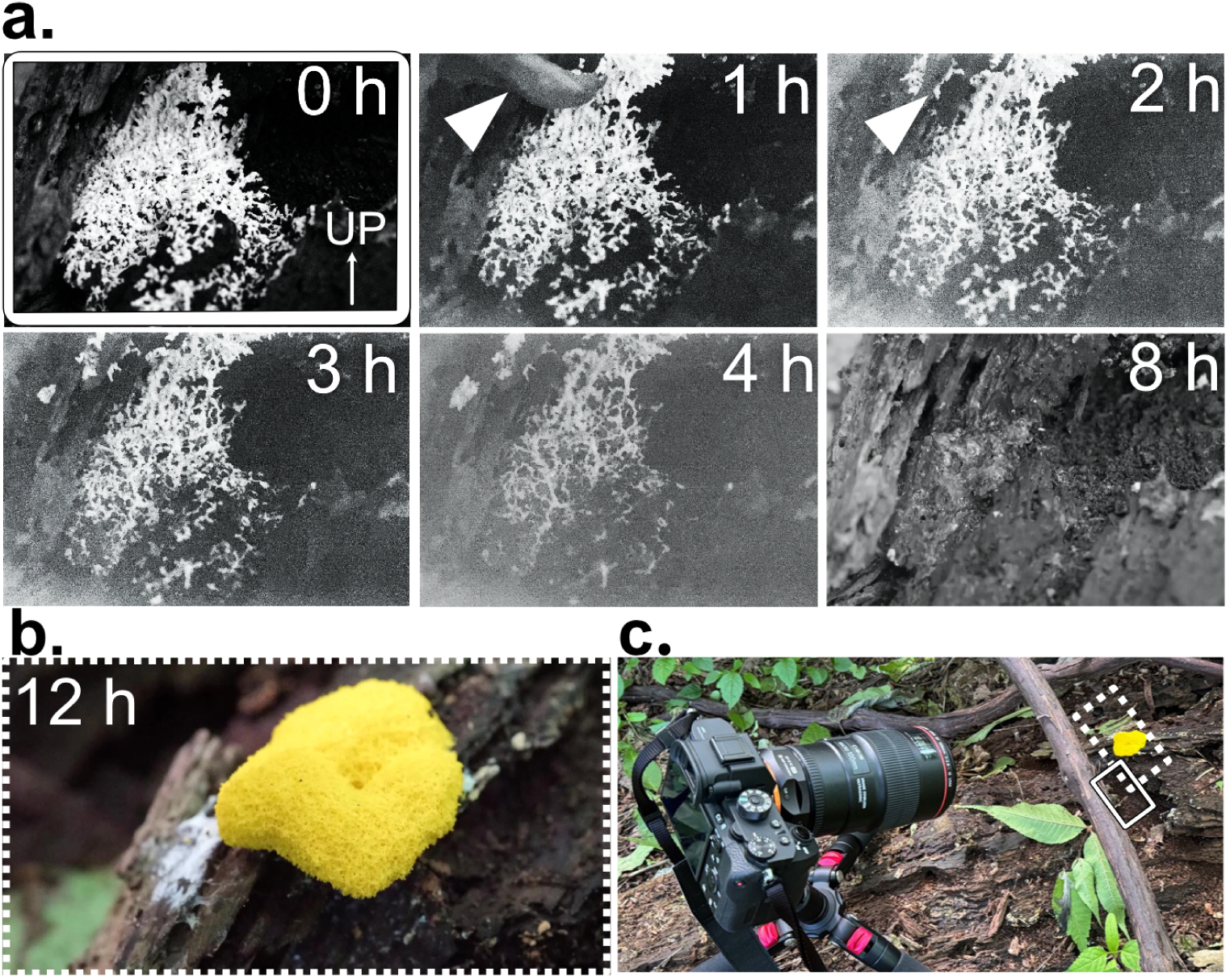
**a**. Climbing behavior of *Fuligo septica* before the sporulation process on a natural fallen tree. Recording started at 18:00 on July 8, 2024 at Hokkaido University (Sapporo). The moving front (giant pseudopod) formed on the uppermost part of the plasmodium (white arrowhead) and led the entire cell to climb over the following hours. At approximately 19:00, a slug visited the plasmodium; the plasmodium (Arrowhead on 1 h) was separated into some pieces after being crawled over by the slug (Arrowhead on 2h). After 8 h, the whole cell went out of the frame; and a fruiting body (FB) was found on the top of the fallen tree. **b**. The FB formed after 12 h. **c**. Wide view of the recording field. The solid line indicates the field of view in a; the dashed line indicates the field of view in b.

### 2.2 Experiments

#### 2.2.1 Organism and culture conditions

The plasmodium of *Physarum polycephalum* was cultured in a rectangular container (24 cm*×*34 cm) on 1% w/v agar plates (agar powder for cooking, type S-7, Ina Food Industries in Ina city, Nagano, Japan) with 0.5 cm thickness in the dark at 25 °C. To prepare the agar plates, 1% w/v of molten agar made from the agar powder was poured into a plastic container. The plasmodium was fed with oat flakes (Quaker Oats Co., USA) twice daily. A part of the agar plate (approximately 1/3 in area of the entire plate) across which the plasmodium was spreading was transferred to a newly prepared agar plate every two days to maintain the stock cultures.

Before the experiment, the agar plate on which the plasmodia crawled and fed on oat flakes was cut out, and placed on a new plain agar plate. Prior to the experiment, the organisms were allowed to crawl freely on an agar plate for up to 24 h without feeding.

Throughout the laboratory experiments, we used a subdivided portion of a single wild-type strain (called *Sonobe* strain hereafter), provided by Prof. Seiji Sonobe at University of Hyogo.

#### 2.2.2 Experiment 1: Effects of undulating surface of (experimental arena) on FB formation

We conducted an experiment to investigate the effect of an uneven surface on sporulation in *P. polycephalum*. We prepared chambers with three types of scaffold (Fig. 3) composed of poly lactic acid (PLA) filament fabricated using a 3D printer (X1E, Bambu Lab, China; see Supplemental Information for printable 3D data) and agar gel. The “flat” scaffold had no uneven surface, whereas the “wave” scaffolds had wave-like structures to create uneven surfaces. We prepared two types of wave scaffolds that were line-symmetric with respect to the axis of the rotating body: Wave I and Wave II (Fig. 3).

**Fig. 3.**
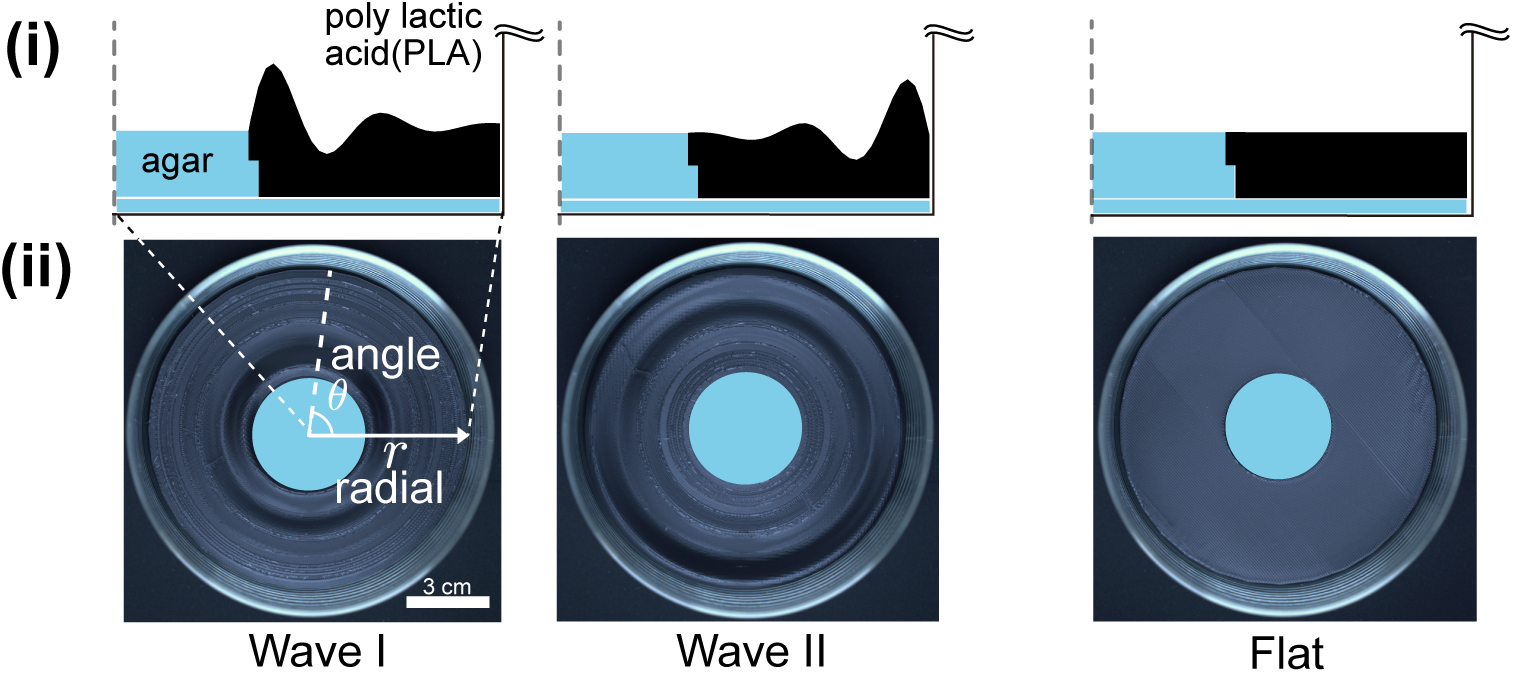
Experimental chamber used in Experiment 1. (i) Schematic illustration of a cross-sectional view. The chamber was designed to be rotationally symmetric. The dashed line (gray) represents the symmetry axis. The resulting revolution was reproduced using a 3D printer and a PLA filament (black) and set into a glass Petri dish with agar gel (blue). (ii) Top view. We defined the radial axis *r* and angular axis *θ* on the top projection of the scaffold.

**Fig. 4.**
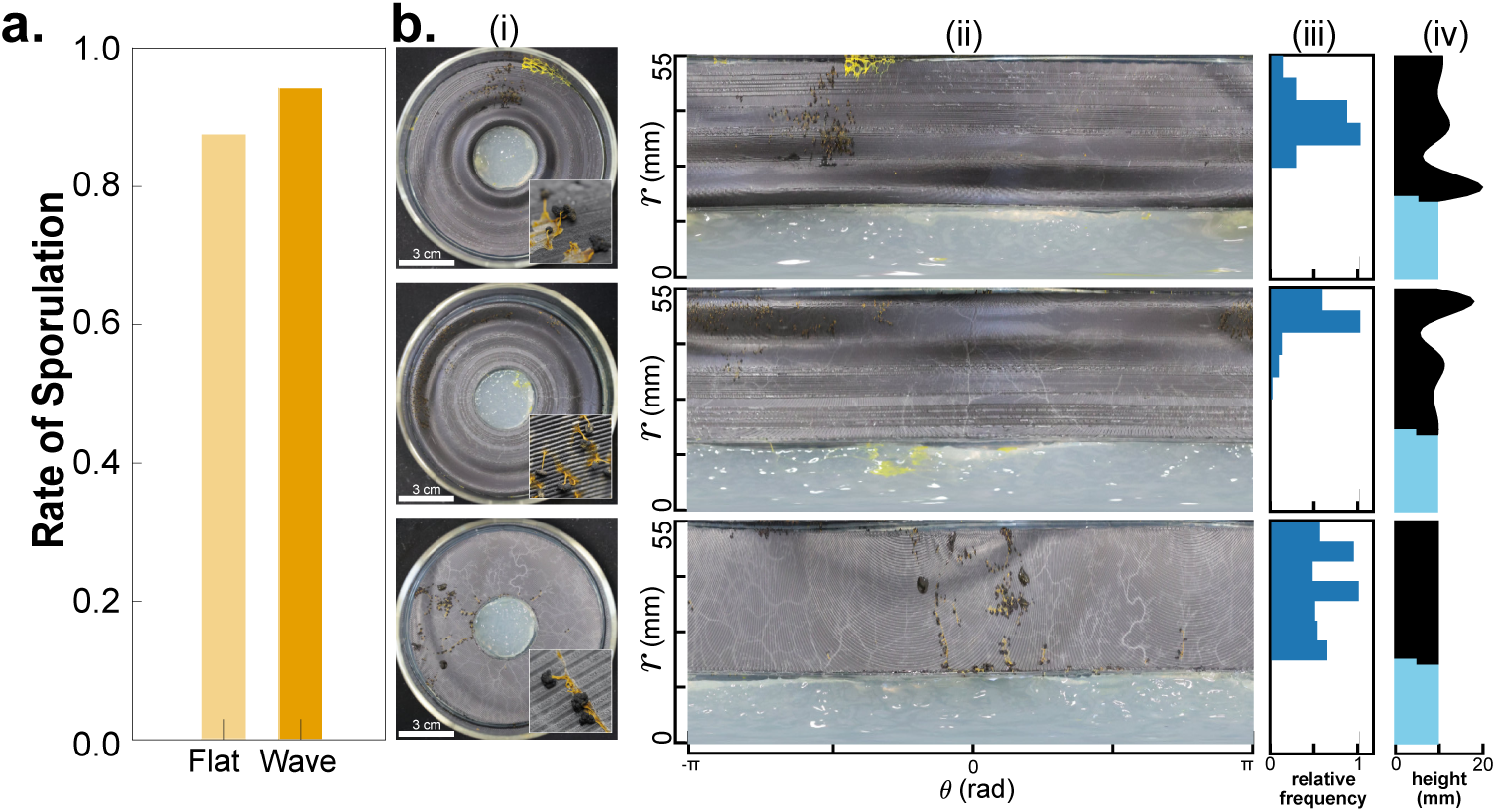
Results of Experiment 1.**a**. Statistical results. High sporulation rate was observed in both types of scaffolds (Flat: 14 out of 16, Wave-I: 9 out of 9, Wave-II: 7 out of 8 samples). **b**. Representative results of the sporulation.Top: Wave-I, middle: Wave-II, bottom: flat-type scaffolds.(i) Top view of the experimental chamber in the final state.Inset: Close-up view.(ii)-(iv) The ordinates within each row, corresponding to the radial direction *r* of the crawling ground, as indicated in Fig. 3(ii).(ii) Results of the polar transformation of the images shown in (i).Abscissa: Angular direction *θ* of the original image.(iii) Relative frequency of fruiting bodies (FBs) in the radial direction.Abscissa: relative frequency of the FBs.(iv) Height modulation of the crawling ground surface.Abscissa: height of the crawling surface.All FBs were formed on the scaffold, particularly on elevated areas when present.

We trimmed the agar plates on which the moving front of the one-day-starved plasmodium crawled to fit into a *ϕ* 35 mm Petri dish(Violamo, AS ONE, Japan). The dish was mounted at the center of a 3D-printed scaffold after the plasmodium had spread densely over the agar surface. The entire setup was enclosed in a glass container (120 mm diameter, 60 mm height, 2.73 *±* 0.50 mm thickness) with a lid. To maintain the humidity, 50 mL of 1% w/v molten agar was poured into the container and allowed to solidify at the bottom before being placed in the setup.

To provide ultraviolet illumination, which has been confirmed to be an effective sporulation stimulus (Nakagaki et al. 1996), a germicidal lamp (GL15, TOSHIBA Tokyo, Japan) was used, with the light positioned 60 cm above the sample and passing through a glass container. Germicidal wavelengths shorter than 350 nm were partially blocked by the glass, allowing minor spectral components, including ultraviolet light of approximately 366 nm, to pass through with an intensity of 17 *µ*mol*/*m^2^*/*s (GL Spectis 5.0, GL Optics, USA; see Supplemental Figure 1 for detailed spectral data).

During the 72-h exposure period, the organisms were allowed to crawl freely on both the scaffold and agar gel. After ultraviolet exposure, the sporulation state of the plasmodia was determined by visual inspection.

#### 2.2.3 Experiment 2: Effects of body size, dry surface, and various elevations in the experimental arena

We conducted another experiment to observe the behavior of *P. polycephalum* during sporulation depending on cell size.

After the plasmodium was starved for one day and spread densely over the agar surface, a small piece of plasmodium was cut from the tip of the starved organism. A plastic film was carefully inserted between the cut piece of the plasmodium and surface of the agar. The film with the slime mold piece was placed on the pan of an electronic balance(BCE2202I-1SJP, Sartorius, Germany); the weights of the film and the plasmodial piece were measured. Subsequently, the weight of the film alone was measured to determine the weight of the plasmodial piece.

The plasmodia were placed on an agar plate 1 cm in front of the acrylic board and allowed to crawl freely for 48 h. The agar plate was placed on a relatively large polystyrene dish to prevent the plasmodia from crawling out of the experimental system. As shown in Fig. 5, we presented three types of acrylic boards placed vertically on an agar gel (010-08725, Wako Pure Chemical Industries, Japan), with different heights (25, 50, and 100 mm), 2 mm thickness, and 20 mm width. The surface areas at three different locations in the experimental arena measured 74,500 mm^2^ (base location: plastic dry surface), 10,000 mm^2^ (agar location at lower elevation), and 7,800 mm^2^ (higher location: acrylic upright board).

During the crawling period, the entire system was enclosed in a cell incubator (LTE-500, TOKYO RIKAKIKAI, Japan) maintained above 95% of relative humidity, and illuminated with collimated ultraviolet light (Hololight, PiPhotonics, Japan) at a 45*^◦^* angle, with a peak wavelength of 378 nm and intensity of 82 *µ*mol*/*m^2^*/*s (GL Spectis 5.0, GL Optics, USA; see Supplemental Figure 1 for the raw spectrum data). Crawling behavior was recorded using a digital camera (Tough TG-7; OM Digital Solutions, Japan). Images were acquired continuously for 48 h at a rate of one frame per 10 min. Based on the acquired image sequence, we manually tracked the highest point of the plasmodial body on each board and converted it into the timeline *h*(*t*) as shown on Fig. 6. After the crawling period, images of the entire dish were obtained to count and locate the FBs. The maximum height of each board was tracked manually. Twenty samples with different wet weights were used.

**Fig. 5.**
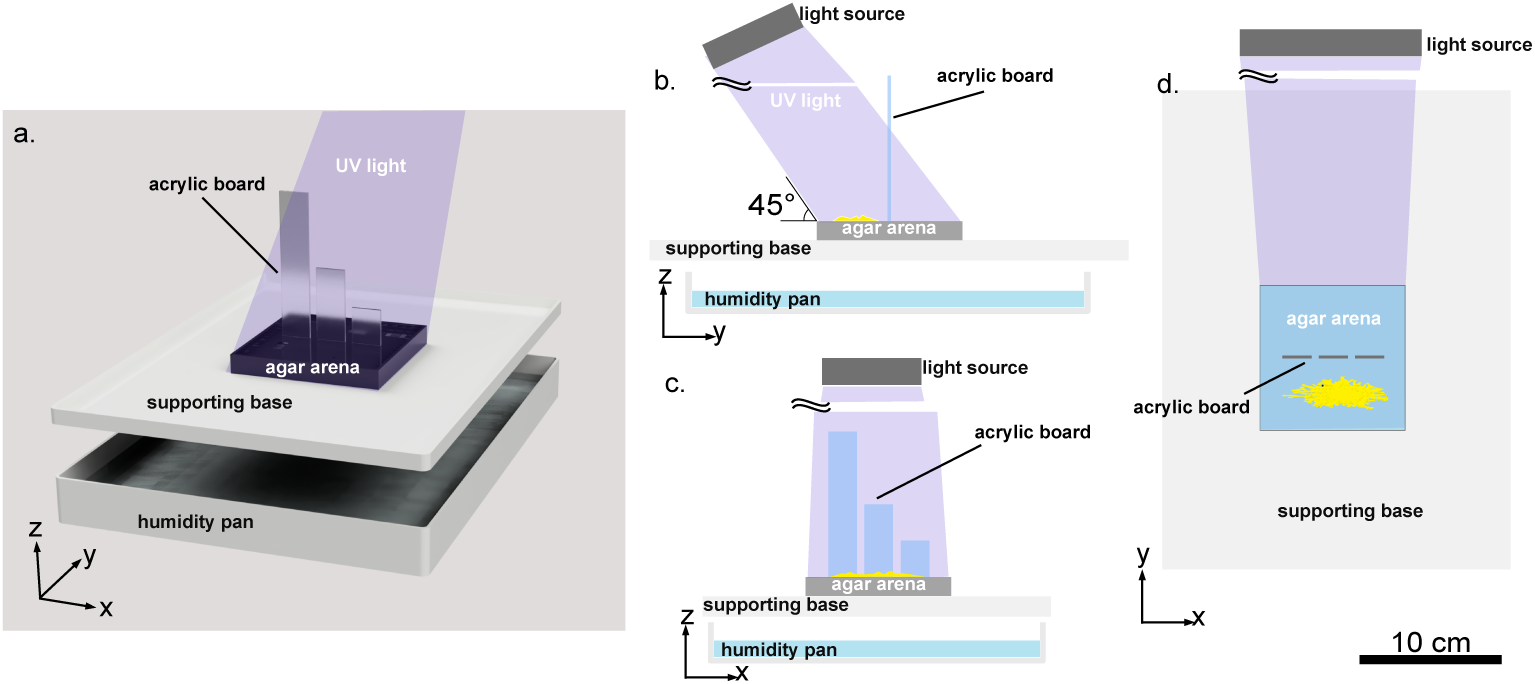
Schematic illustrations of the experimental chamber used in Experiment 2. **a**. Bird’s eye view, **b**. Side view, **c**. Front view, **d**. Top view. The scale bar on the lower right is for b, c, d.

**Fig. 6.**
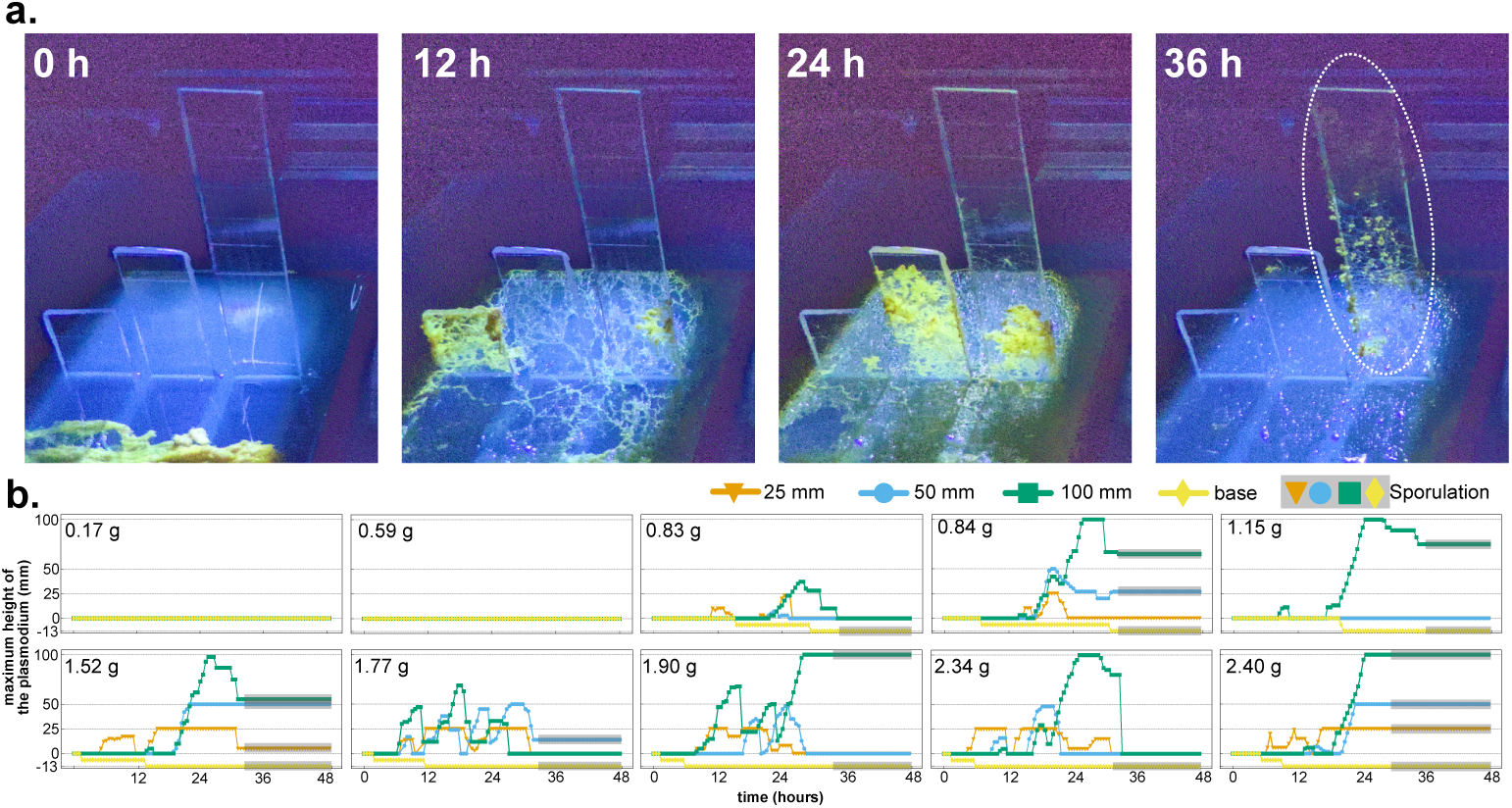
**a**. Example sequence of the recorded images. The wet weight of the sample was 1.90 g. The dashed line indicates the segmentation of the plasmodium, which we define as the initiation of the sporulation process. **b**. Ten typical exploration processes over 48 h on acrylic boards of 25 mm (orange), 50 mm (blue), and 100 mm (green) height. The maximum height of the plasmodium crawling on each board was plotted. For reference, we also plotted the minimum height, those lying on the dish, and the base area (outside of the dish), as indicated in yellow. Abscissa: time (h), ordinate: height (mm) in each panel. The dashed gray horizontal line indicates the top height of each board. Shaded markers indicate the occurrence of sporulation in the area.

#### 2.2.4 Data analysis

For the analysis of the results in Experiment 1, we defined a polar coordinate system over the top aspect of the circular crawling ground (Fig. 3(ii)) and unwrapped it into a rectangular view to identify the radial location of FBs on uneven circular surfaces. Additionally, we compared the sporulation rates between undulating (Wave I:*N* = 9, Wave II:*N* = 8) and flat (*N* = 16) scaffolds using a two-sided Fisher’s exact test.

In Experiment 2, we systematically varied the wet weight of the plasmodium across the experiments. We assessed regression slopes for wet weight versus FB height, crawling height, and FB number and ratio (Fig. 7 and Fig. 8-f).

**Fig. 7.**
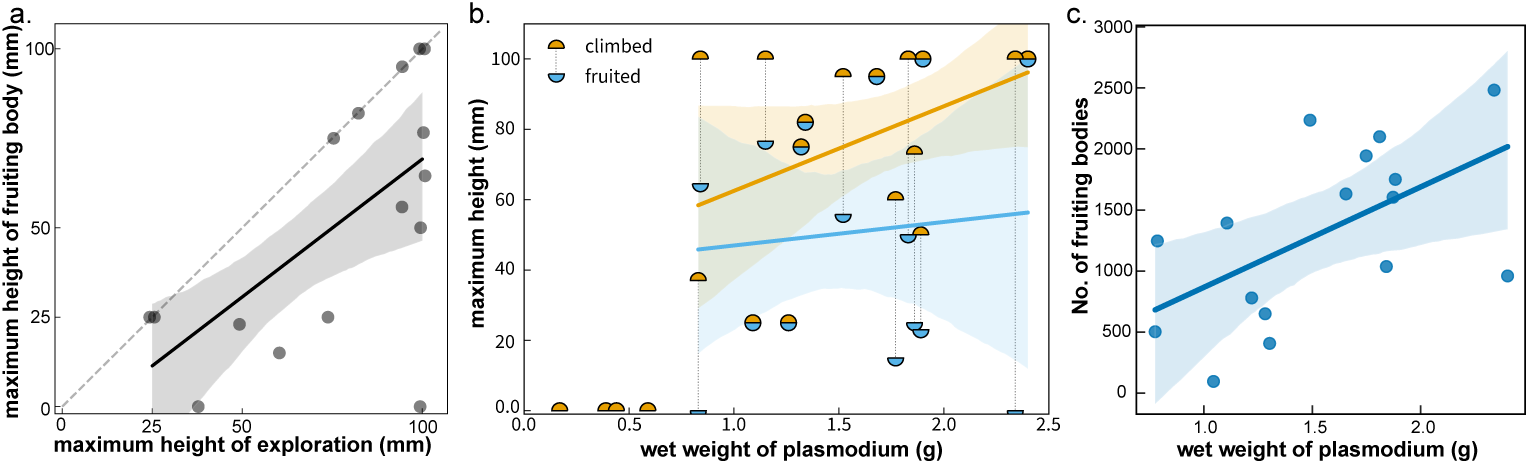
**a**. The maximum height reached while exploring (abscissa) vs. the maximum height of fruiting body (FB) formation (ordinate), plotted with gray solid circles for 20 samples, compared with a dashed line indicating the limit of the FB height. The solid line and the translucent area indicate the linear regression line and its 95% confidence interval, respectively. **b**. The relationship between biomass (abscissa) and the same two variables as in (a) (ordinate, orange:climbed, blue:fruited). Both variables had a low correlation with biomass. The solid line and translucent area indicate the linear regression line and its 95% confidence interval, respectively, considering data that exceed the minimum fruiting weight (0.83 g). **c**. Total number of FBs vs. wet weight of plasmodium. The circular dots represent observed data points. The solid line and surrounding pale blue area represent the linear regression line and its 95% confidence interval, respectively. As the regression line shows, approximately 1 g plasmodium is expected to form 750 FBs. The maximum weight of a non-sporulating plasmodium (0.59 g) was predicted to be sufficient to produce 400 FBs.

**Fig. 8.**
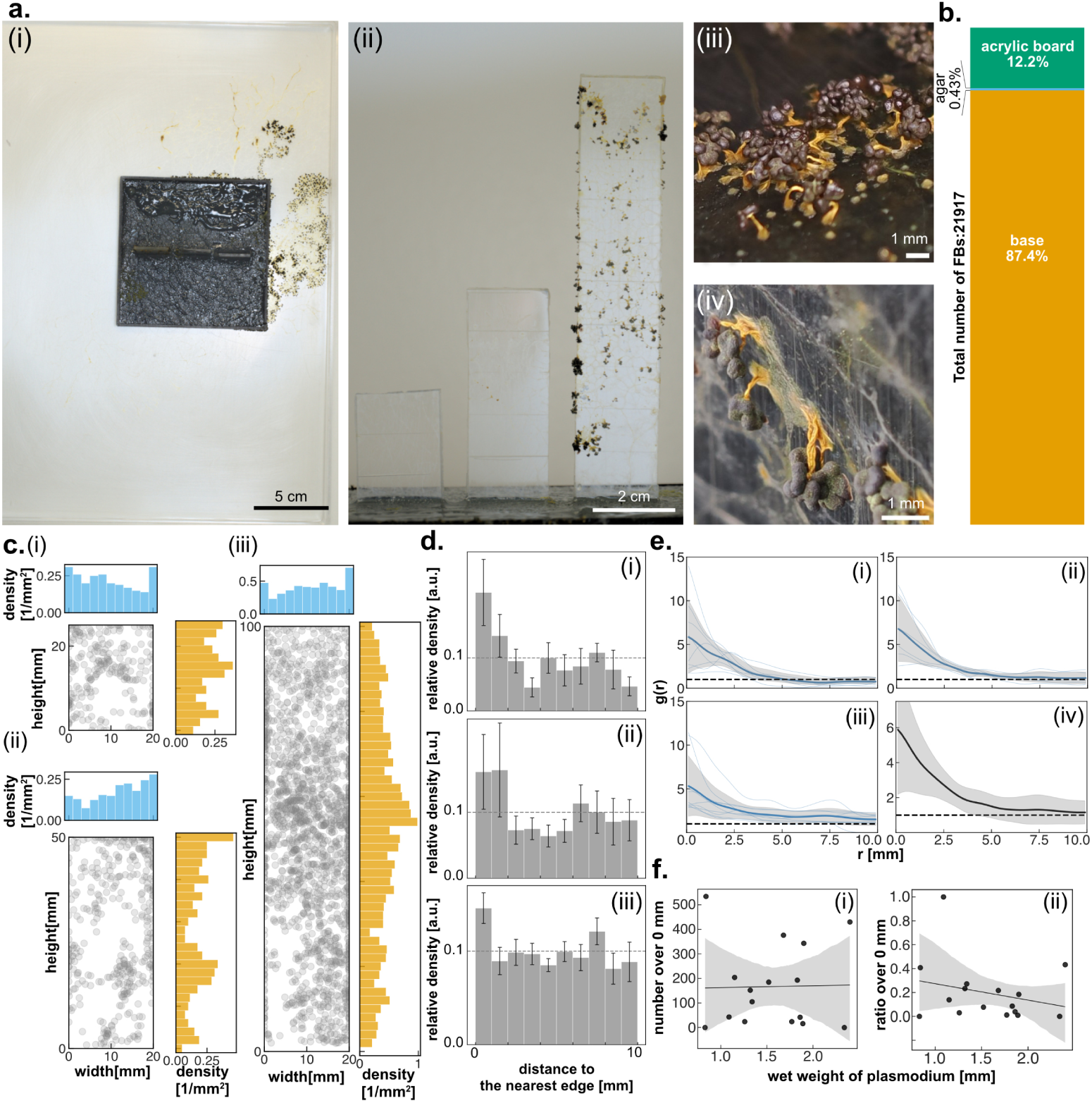
Distribution of fruiting bodies (FBs) in Experiment 2. **a**. Representative results for fruiting. The sample is the same as the one shown in Fig. 6-a. (i) Top; (ii) lateral view of the experimental system; (iii) and (iv) close-up view of the base area (iii) and 100 mm acrylic board (iv). **b**. Total number of FBs obtained from all 20 plasmodia and its location ratio. **c**. Spatial distribution of FBs on each acrylic board. (Scatter plot, gray): Distribution of counted FBs on each acrylic board. (Histogram): Number density of FBs along *x* and *y* axes. (i), (ii), (iii) correspond to the result in the 25, 50, 100 mm boards. **d**. Histogram of FB density categorized by the nearest distance from the edge. (i), (ii), (iii) correspond to the result in the 25, 50, 100 mm boards. **e**. The radial distribution function *g*(*x*) calculated from (i) 25 mm, (ii) 50 mm, (iii) 100 mm, and (iv) overall distribution. Dim gray envelope indicates the standard deviation. **f**. Number (i) and ratio (ii) of the FBs located on the acrylic board. The solid lines and translucent areas indicate the linear regression line and its 95% confidence interval, respectively.

We estimated the threshold body size for FB formation as the midpoint between the largest size of the plasmodium at which FBs were not formed and the smallest size at which FBs were formed. To assess the uncertainty of this estimate, we applied a bootstrap procedure by resampling 100,000 times from the datasets, allowing duplication; a 95% confidence interval was yielded.

We counted and recorded the position of all FBs across the experimental arena. For each board, we superimposed the distributions of FBs across all experiments and visualized them in Fig. 8-c. Since the board has a finite thickness, an FB near the edge can lie on the front face, the back face, or the 2 mm-thick side face. To keep the effective bin width along the surface constant, the bin width was set to 1 mm at the edge, and 2 mm in the interior of each board. At the edge, a bin spans 1 mm on the front face, 2 mm on the side face, and 1 mm on the back face, giving an effective width of 1+2+1=4 mm; in the interior of the board, a bin spans 2 mm on each of the front and back faces, giving 2+2=4 mm. For the height direction of the 25 mm board only, the bin width was set to 1.92 mm in the interior area not to yield a uneven bin width, giving an effective width of 3.83 mm.

The relative number density was also calculated and binned based on each FB’s distance from the edge of each board (Fig. 8-d). The procedure is as follows. For each board in each experiment, a rectangular region of interest was defined with height *h*_max_ = max*{h*(*t*)*}* and width 20 mm. Then, offset lines were drawn at 1 mm intervals from the edge. The area of each 1 mm-wide band was computed, and the number of FBs within each band was counted. The number density was calculated as the count of FBs divided by the band area. Finally, the number density of each bin was normalized by the sum of densities across all bins to obtain the relative density, which represents the probability of finding an FB in each band. The difference between the resulting relative number density and the relative density expected under a uniform distribution was tested across experiments using two-sided Wilcoxon signed-rank test.

To study statistical distributions of the distances among FB, we used the radial distribution function, which is typically used to quantify the radial distribution of the number of particles around a particle. In our analysis, it was used to quantify the radial distribution of the number of FBs around an FB. The geodesic distance *r* between every pair of FBs on the upright board was computed; the number density of FB was calculated as *p*_obs_(*r*). *p*_obs_(*r*) was averaged over each board to obtain *p*^_obs_(*r*). A smooth *p*^_obs_(*r*) estimate was obtained via Gaussian kernel density estimation with a bandwidth of 0.5 mm. To normalize against a random reference, we randomly placed the same number of FBs uniformly on each face of the board 999 times numerically and averaged across trials to obtain *p*^_sim_(*r*). The radial distribution function is defined as *g*(*r*) = *p*^(*r*)_obs_*/p*^_sim_(*r*). Hence, *r*_cluster_, at which *g*(*r*) first crosses *g*(*r*) = 1, indicates the characteristic cluster radius, beyond which *p*^_obs_ becomes random or repulsive to each other rather than attractive. For calculations to obtain *g*(*r*), the crawled area of the upper board was approximated as two flat faces (front and back), each with area (*h*_max_ *×* 20) mm^2^, neglecting the side surfaces(See Supplemental Figure 2).

We used the multi-point tool in Fiji version 1.54f (Schindelin et al. 2012) to count and locate FBs, OpenCV 4.10.0 (Bradski 2000) to conduct polar coordinate expansion, Scipy 1.14.0 (Virtanen et al. 2020) to conduct Fisher’s exact test, Wilcoxon signed-rank test, and kernel density estimation, and seaborn v.0.13.2 (Waskom 2021) and statsmodels 0.14.6 to generate regression plots.

## 3 Results

### 3.1 Field observations for two species, *Physarum rigidum* and *Fuligo septica*

#### 3.1.1 Crawling around a decaying log and producing FBs multiple times while changing location, observed in *Physarum rigidum*

Fig. 1-a presents an overview of the observation site on the forest floor. The plasmodium of *Physarum rigidum* crawled over the surface of a decaying log on the forest floor for seven days. Fig. 1-b presents a schematic summary of the time course. The plasmodium did not simultaneously form FBs. Instead, part of the plasmodial body transited into FBs four times at different locations.

We plotted this trajectory of crawling every 2 h on the *x*-*z* coordinate determined arbitrarily, where *x* and *z*-axes correspond to the horizontal and vertical locations, respectively (Fig. 1-c). The color of the data points indicates the time sequence from the beginning of the recording(violet at the beginning and yellow at the end of the observation period). The plasmodium migrated from the lower right on June 28, 2025 to the upper left on July 3, 2025, over six days while initiating FB formation three times, at 17:00 on June 29, at 22:00 on June 30, and at 12:00 on July 1. Because the data points are often overplotted, we indicate the frequency of the data points by the histograms (orange bars along the *x*-and *z*-axes, Fig. 1-c). Both *x* and *z* histograms indicate that data points are relatively frequent at sites where FBs are formed, indicating a slow crawling speed.

After three times of sporulation, the plasmodium continued to crawl toward the upper side of the tree within the following three days, finally reaching the topmost part of the tree and sporulating with the largest number of FBs (Fig. 1-b and d). Other organisms of slime mold such as *Lycogala epidendrum* and *Arcyria cinerea* also formed FBs at this topmost location. Patches of fruiting bodies were often visited by other species (Fig. 1-e) such as isopods (Fig. 1-e(i)) and insects (Fig. 1-e(ii) and (iii)).

In the field, over the course of one week, the plasmodium moved along the side of a fallen tree for nearly 1 m and formed FBs in multiple locations, including dark and damp areas near the base of the tree and bright dry areas near the top.

The spatial distribution of the FBs is shown in Fig. 1-b. The numbers indicated by the yellow ellipses in Fig. 1-b show the number of FBs formed at this location. The date and time of sporulation were also indicated, estimated from the time the plasmodium separated into segments that were transformed into FBs. As indicated, the plasmodium crawled across fallen trees for a week; 61.1% of FBs formed in the topmost part of the tree. In total, 38.9% of the FBs were formed at the lower part of the tree.

#### 3.1.2 Crawling upward and simultaneously forming FBs at the same location, observed in *Fuligo septica*

In the case of *Fuligo septica*, we observed a specific climbing behavior before sporulation began. After crawling through and exiting the frame for 8 h (Fig. 2-a), the plasmodium formed FBs on the upper part of the fallen tree within 4 h (Fig. 2-b). The field of view is shown in Fig. 2; a and b are indicated by white solid and dotted lines, respectively, in Fig. 2-c.

The plasmodium crawled upward, and a whole plasmodial body simultaneously became FBs at the same location. Additionally, the plasmodial body was visited by a slug during the crawling period and separated after the visit (Fig. 2-a).

### 3.2 Experiment 1: Biased allocation of FBs to the undulating surface of experimental arena, observed in *Physarum polycephalum*

In Experiment 1, we investigated the effects of undulating surfaces on sporulation in *Physarum polycephalum*. After 72 h, over 85% of the samples exhibited sporulation on the wave and flat scaffolds. As shown in Fig. 4-a, no significant effect of surface modulation on the sporulation rate was detected (*p* = 0.6012, two-sided Fisher’s exact test).

Fig. 4-b (i) shows typical results for the final states of each type of scaffold. To observe where the FBs were formed, the circular arena was converted into a rectangular shape through polar transformation, as shown in Fig. 4-b (iii). It was clear that the FBs were formed on the scaffold surface rather than on the agar surface; the relative frequency of sporulation locations was slightly related to the elevation of the scaffold surface in the undulating arena, although there was some deviation in the relative frequency of sporulation locations, even on the flat surface. The spatial distribution of FBs was more biased in the undulating scaffold than in the even scaffold (Fig. 4-b (iii) and (iv)). For the undulating scaffold, locations with high relative frequency were at higher elevations within the undulating arena.

### 3.3 Experiment 2: Location preference of FB formation for a dry and elevated place in relation to the body size of *Physarum polycephalum*

#### 3.3.1 Crawling trajectory before FB formation

We tested the effects of body size, physical nature of the crawling surface, and elevation of the arena on FB formation. We obtained the crawling trajectory of plasmodia on acrylic boards over 48 h by analyzing the image sequence (Fig. 6-b).

Relatively small plasmodia with 0.17 and 0.59 g of wet body weight crawled around only on the wet agar plate and did not move out from the agar plate into the dry surfaces either at the lower (−13 mm height from the agar plate) or higher elevation on the upright boards. The threshold size was estimated as 0.71 g, as a midpoint between 0.59 g and 0.83 g, with 95% confidence interval of [0.61, 0.87] g by a bootstrap procedure (see Methods for detail).

Plasmodium larger than 0.83 g climbed upright boards and often moved up and down across them, resulting in a complex trajectory at elevations (Fig. 6-a and b). While climbing up the boards, it also explored the dry surface at the lower elevation (−13 mm).

When being larger than 0.83 g, plasmodia often reached the highest position on the boards (100 mm height); however, they did not always remain there. Certain samples moved down the middle height of the upright boards or the bottom of the arena, as shown in the timeline of the 2.34 g sample(Fig. 6-b).

#### 3.3.2 Timings and locations of FB formation in relation to crawling behavior

From all 20 tested samples, 16 plasmodia that exceeded the threshold body size mentioned in the previous section formed FBs within 48 h of crawling. Here, we defined the initiation of FB formation as the onset of a morphological transformation specific to FB formation (specific segmentation of the plasmodium into small pieces, each of which is a single FB). Fig. 6-b shows the timing and location of FB formation onset in relation to the crawling trajectory, as indicated by the black symbols. FB formation occurred simultaneously, even when the plasmodial body was spread across different heights.

The remaining four plasmodia below the threshold body size continued to crawl on the wet agar surface without forming FBs.

#### 3.3.3 Relationship between two maximum elevations to which the plasmodia climbed up and formed FBs

The maximum elevation at which the plasmodia crawled correlated with the maximum elevation at which they formed FBs (Fig. 7-a).

Figure 7-b shows the relationship between the weight of plasmodium and the maximum elevation at which it crawls (orange symbols) or forms FBs (blue symbols). The weight threshold for crawling behavior (0.59–0.83 g) coincided with the threshold for FB formation. Above this threshold, the plasmodia exhibited climbing behavior and formed FBs at even higher elevations.

#### 3.3.4 Numbers and locations of FBs formed after crawling

Fig. 7-c shows the relationship between the weight of plasmodium (limited to values above the threshold) and the number of FBs formed. The number of FBs increased proportionally with the weight of the plasmodium. This relationship indicates that even a plasmodium at the threshold weight is sufficiently large to form approximately 400 FBs. Therefore, plasmodia below the threshold were not incapable of forming FBs; rather, under these experimental conditions, certain factors prevented the formation of FBs.

Data that deviated from this proportional relationship were occasionally observed. For example, a plasmodium weighing 1.09 g produced only a small number (44) of FBs. This is because not all parts of the plasmodium became FBs; only a portion did. The phenomenon in which only a portion of the body mass formed FBs while the rest of the plasmodium continued to crawl was also observed in field observations of *Physarum rigidum*. The plasmodium of *Physarum polycephalum* used in the present study mostly produced FBs from all segments simultaneously, whereas at other times, only a portion did so. However, the cause of this variation remains unclear.

#### 3.3.5 Spatial distribution of FBs in the experimental arena

The FBs formed at higher places on the upright boards were distributed in a patchy manner(Fig. 8-a(i), (ii)). Each FB consisted of a yellow stalk(*∼*1 mm in length) and black sporangium(*∼*1 mm in diameter) on its top, as shown in Fig. 8-a(iii), (iv). Over 87.4% of the FBs formed at the lower places (−13 mm elevation) rather than on the agar area (0 mm elevation) (Fig. 8-b). The remaining 12.2% formed on areas higher than the agar area. Only 0.43% of the FBs formed on the agar area (Fig. 8-b).

The location of FBs were overlaid within each board (*N* = 9 at 100 mm board, *N* = 8 at 50 mm board, *N* = 6 at 25 mm board) and shown in Fig. 8-c. with histograms of number density along height and width direction. We observed a high density in the edge area of each board except for the top edge of the 100 mm board (Fig. 8-c(i)). FBs were concentrated on the middle-elevation area on the 100 mm board instead of on the top edge.

Fig. 8-d shows the relative density of FBs as a function of the distance from the nearest edge, which was calculated across 25, 50, and 100 mm boards (i, ii, iii) and averaged across the experiments (iv). Gray dashed lines in each panel indicate the theoretically expected value of relative density for a uniform distribution (0.1). The relative density in each board in each experiment was compared to the the value 0.1 using the two-sided Wilcoxon signed-rank test. Only the first bin, namely the area within 1 mm of the edge, showed a significant difference (*p <* 0.05) with positive deviations from the uniform distribution.

The radial distribution function normalized against the uniform distribution *g*(*r*) is shown in Fig. 8-e. All *g*(*r*) have a *g*(*r*) *>* 1 region, showing the existence of a cluster; the curve rapidly decreases at approximately *r* = 5 mm. However, except for the 25 mm board (Fig. 8-e(i)), we could not observe the characteristic cluster size *r*_cluster_ which is obtained as the first intersection against *g*(*r*) = 1 on average (Fig. 8-e(ii)-(iv)). See Supplemental Data 2 for the detailed distribution in each individual.

The number of FBs at the upper boards ranged from 200 to 600 in samples observed in this study; no clear dependence on body weight was shown (Fig. 8-f(i)). Conversely, the ratio of FBs at higher locations showed a decreasing manner to the body weight(Fig. 8-f(ii)).

## 4 Discussion

### Size-threshold for sporulation and its prior dry-surface exploration

Throughout the laboratory experiment using *P. polycephalum*, we rarely observed FBs forming on wet agar (Figs. 4-b, 8-b). Moreover, plasmodia did not form FBs unless they crawled over a dry surface (Figs. 6-b, 7-b). This finding may be consistent with the behavioral changes before sporulation in certain species; taxis toward dry surfaces switches from negative to positive immediately before sporulation (Stahl 1884; Lubbock 1892; Constantineanu 1906).

In Experiment 2, we observed the size threshold at which plasmodia initiated migration away from the agar(Fig. 7-b). Importantly, this threshold coincided with the onset of the transition to FB. All plasmodia below the threshold (*N* = 4) formed no FBs, whereas all plasmodia above it (*N* = 16) transformed their entire body mass into FBs, with the exception that only a part of the body was transformed into FBs (1.09 g). Consequently, the number of FBs and wet weight of the cells showed a linear relationship (Fig. 7-c). According to an estimation from this relationship, the critical body size was sufficient to yield approximately 400 FBs.

On the behavioral strategy for reproduction in multicellular organisms, Gadgil and Bossert (1970) showed that the big-bang reproductive strategy was optimal under certain conditions. Here, we discuss the applicability of this theory to FB formation in slime molds. They suppose the reproductive effort *θ*, which remains zero over life stages and rises to the suicidal value *θ* = 1 when the reproductive stage begins in the big-bang strategy. Optimality of reproduction holds when the profit function (the relationship between one’s reproductive effort and fitness gain in its offsprings) is convex (convex downward) or when the cost function (the loss in future reproductive potential associated with current effort) is concave (convex upward), assuming that both functions are monotonically increasing.

It is reasonable to assume that in slime molds, the reproductive effort *θ* is proportional to the number of spores *N* . The probability that one spore out of *N* spores survives is expressed as a concave function of *θ*: *P*_simple_ = 1 *−* (1 *− p_s_*)*^N^*, where *p_s_* is the per-spore survival probability. Here, we introduce the feature of the life cycle of slime molds: a plasmodium is diploid and develops after two haploid amoebae with a specific mating type germinate from spores and mate each other. This means that multiple amoebae need to co-arrive at the same habitat patch. The probability of such co-arrival is given by 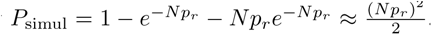 . This estimation was derived from the leading term of the Poisson approximation, in which at least two spores co-arrive in the same niche under the condition *Np_r_ <* 1, where *p_r_* is the probability for a spore to reach a niche. The threshold size for FB formation in slime molds is to be interpreted as an optimal strategy according to the conventional big-bang reproductive theory as the probability *P*_simul_ is a convex function of *θ* and satisfies the condition for the optimality of the big-bang strategy.

One of the above theoretical consequences from the big-bang strategy is that the number of spores is substantially higher in sexual reproduction than in asexual reproduction because the co-arrival probability is substantially lower than the probability of single arrival. In fact, Schnittler and Tesmer (Schnittler and Tesmer 2008) reported that the Myxomycetes, a group of sexually reproducing eumycetozoans including genus *Physarum* and *Fuligo*, produces substantially more spores than asexually reproducing eumycetozoans, such as Dyctiostelids and Protostelids.

The bang-bang strategy, a similar theory to the big-bang strategy, was applied to the collective behavior of an ant colony (Oster and Wilson 1978). The results obtained here in slime molds have suggested a possibility of similar theory in a huge, multinucleate amoeba. The fact that this strategy can be applied to various lifestyles with different life cycles suggests the existence of general traits in reproductive behaviors.

### Effects of available dry surface on sporulation

In Experiment 1, we initiated ultraviolet irradiation of the plasmodia without prior starvation. A high sporulation rate was observed after 72 h of crawling and irradiation (Fig. 4-a). In Experiment 2, we observed the onset of morphogenic changes for sporulation (segmentation) after approximately 36 h, which was completed within 48 h of treatment. Prolonged starvation for *>*72 h (Camp 1937; Wilkins and Reynolds 1979; Chapman and Coote 1982; Akitaya et al. 1985; Starostzik and Marwan 1995a) followed by exposure to ultraviolet or infrared light (Starostzik and Marwan 1994, 1995a; Nakagaki et al. 1996; Marwan and Starostzik 2002) is required for FB formation. Starvation has been recognized as a trigger for pre-morphogenetic changes preceding sporulation, such as gene regulation and metabolic reactions. Subsequent light exposure triggers the onset of morphogenesis within a few hours. The entire process requires approximately 90 h (Glöckner and Marwan 2017), nearly twice as long as that observed in the present study. Previous findings were primarily derived from experiments on uniformly flat and moist agar gels. Therefore, our results on agar avoidance and rapid sporulation indicate that dry surfaces facilitate FB formation by reducing the required starvation period. Furthermore, a recent study of the light and starvation trigger for FB formation among *Physarum* species, including *P. polycephalum* (Masui et al. 2025), reported that the known set of triggers did not work for every species in the genus *Physarum*, suggesting that different pathways or necessary conditions existed. A quantitative evaluation of this effect remains a subject for future research.

### Spatial allocation pattern of FBs

Motivated by the field observations and Experiment 1, we hypothesized that FBs may be formed at elevated positions. To examine this hypothesis, in Experiment 2, we designed an experimental arena with three standing thin boards on a flat agar block and observed FB formation of *P. polycephalum*. In this arena, relatively large plasmodia tended to reach higher elevations (Fig. 7-b) when they freely migrated around there. However, the number of FBs formed at higher elevations was lower than that at lower elevations of the flat area, although there was still a plentiful amount of space available for FB formation on the upper board (Fig. 8-f). Interestingly, regardless of the body size of the organism, approximately 10 to 20% of FBs were formed on the upper boards (Fig. 8-b,f). This result may be related to bet-hedging behavior because higher elevation may have a relatively high risk of FB formation owing to excessive dryness and high efficiency of spore dispersal. This speculation is worth studying because bet-hedging behavior has not been reported in unicellular amoeboid organisms.

Looking at the location of FBs on upper boards, there was a clear tendency that FBs were just on the edges of the board (Fig. 8-c,d). This tendency was also observed in Experiment 1. FBs were more abundant near the top than at the bottom of the undulating substratum(Fig. 4-(iii), (iv)). The ethological meaning of this tendency is unclear; however, it may be related to spore dispersal.

In general, other important factors, such as gravity, surface chemistry and microstructure of the ground substratum, and the microclimate of humidity, light, and wind, may affect FB formation.

Further studies on compound effects of multiple factors on FB formation are highly required in the future so that the potential ethological capacity of slime molds can be explored.

### Issues in studying FB formation in plasmodial slime molds, Myxogastria

Through laboratory experiments with *P. polycephalum*, plasmodia tended to form FBs at multiple locations. We also observed a similar tendency in the field observations in *P. rigidum*, while a plasmodium of *Fuligo septica* moved upward and formed FBs in one location rather than in multiple locations, although each observation was obtained from a single individual of each species. In this species, the FB is formed as a single mass, as reported in the literature (e.g. Stephenson (2025); Manaaki Whenua-Landcare Research (2023)), being called “aethalium” (Leontyev et al. 2019). Compared with many small, separated FBs (stalked sporocarp) in *Physarum* sp., the single large mass of FB in *Fuligo* aethalium may be more tolerant to dry environments.

Schnittler (Schnittler 2001) discussed a “resource allocation model” on FB formation in order to explain selection pressure in the morphology and locations of FBs after extensively observing the morphology and location of FB in many species in the natural field. He suggested two possible strategies for risk-hedging against FB formation (failure, prey, less dispersion of spores, etc.): many small stalked FBs typically observed in *Physarum* species and a single large mass of FB called “puffball strategy” typically observed in *Fuligo* species. This suggestion is consistent with our findings. Our results support this suggestion by designing a well-controlled experimental arena in which certain variations of elevation, ground undulation, and surface character are involved. Further studies on the ethological relationship between morphology and the location of FB formation are required in the future as it is key in elucidating the potential ethological capacity of single-celled organisms.

## Supporting information

Description of Supplemental files and figures

printable stl files

## Acknowledgments

This study was supported by the Sasakawa Scientific Research Grant from the Japan Science Society (ST); additionally, it was supported by MEXT KAKENHI Grant Number JP21H05310 (TN, AT), JP21H05308 (YN) in the program “Ethological dynamics in diorama environment”, Japan Society for the Promotion of Science (JSPS) KAKENHI Grant numbers JP24K09388 (YN) and 26H02509 (TN), the Cooperative Research Program of the Network Joint Research Center for Materials and Devices (YN), the Program for Fostering Researchers for the Next Generation conducted by the Consortium Office for the Fostering of Researchers in Future Generations, Hokkaido University (YN), and the Project of Junior Scientist Promotion at Hokkaido University (YN). The authors deeply thank the reviewers and editors who provided intensive discussions on the ecological aspects.

## Data availability

The datasets generated and analyzed in the current study are available from the corresponding author on reasonable request.

## Compliance with Ethical Standards

The authors declare that this study was conducted in accordance with the ethical standards.

## Ethical approval

We used laboratory-maintained slime molds in experiments; thus, no permits were required for this research. All research was conducted in accordance with the laws and regulations of Japan and Hokkaido University.

## Informed consent

Not applicable to this study.

