## Supplementary material for "Allocation pattern of fruiting bodies in plasmodial slime molds, and threshold size for sporulation of *Physarum polycephalum*": Description of Supplemental files and figures

#### 1 Supplemental materials

- printable 3D Datas
  - **sup_flat.stl**: printable data for "Flat" type scaffold.
  - **sup_inc.stl**: printable data for "Wave-I" type scaffold.
  - **sup_dec.stl**: printable data for "Wave-II" type scaffold.

### 2 Figures

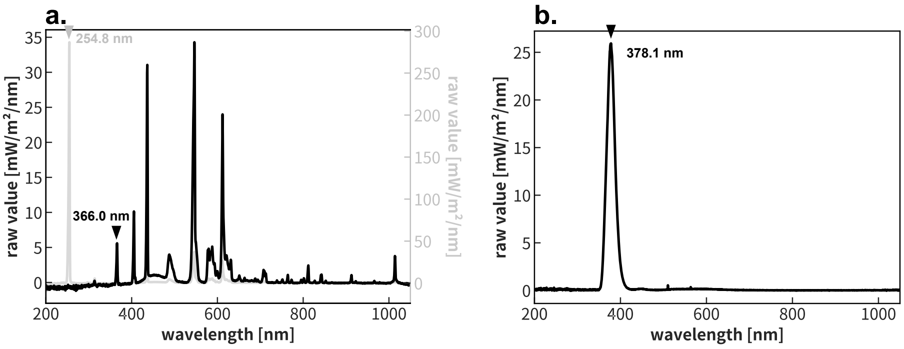


**Fig. 1** Wavelength spectrum of the light source used in experiments. Peak ultraviolet wavelength was marked with arrowhead and the value of wavelength was noted. **a**. Spectrum data of germicidal light used in Experiment 1. Black: inside of the grass container, Gray: direct illumination. **b**. Spectrum data of on the agar area of the arena used in Experiment 2.

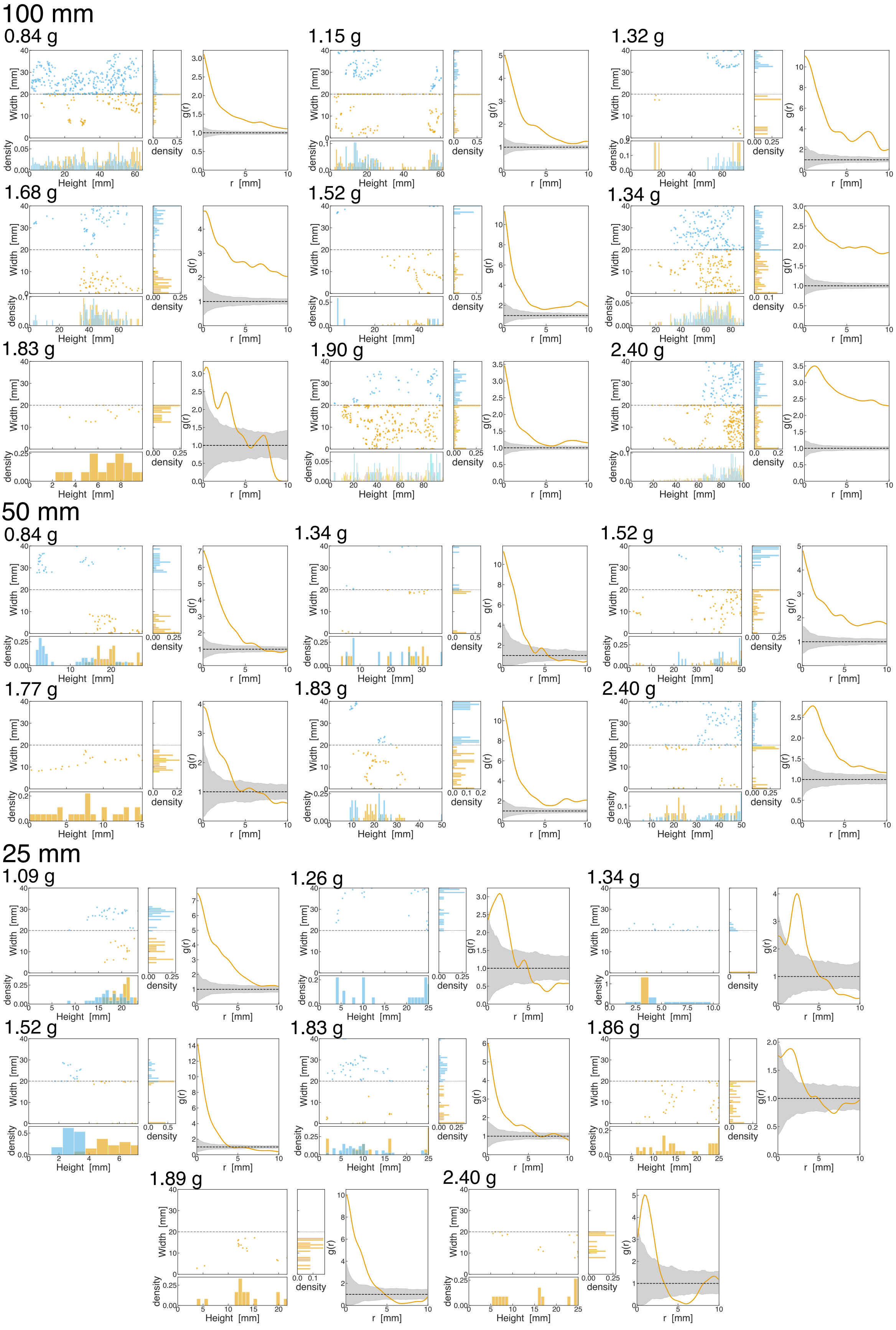


**Fig. 2** Spore distribution on upright board in every experimental replicate. (left) Scatter plot: the raw distribution of the FB location on the board. histograms: relative density of FBs along the axis. (right) solid line: Radial distribution function $g(r)$ derived by following the method indicated in section 2.2.4 in the main text. Gray envelope: 95% confidence limit of homogeneity, derived from the numerical simulation.
